# Comprehensive bioinformatic analysis reveals novel potential diagnostic biomarkers associated with monocytes in osteoporosis

**DOI:** 10.64898/2026.03.20.713320

**Authors:** Xianyun Qin, Bo Wen, Pan He, Zetong Chen, Shengyun Tan, Zhiyou Mao

## Abstract

Osteoporosis affects millions of women globally. In this study, we applied bioinformatics methods to screen for novel diagnostic biomarkers of osteoporosis in women using the GSE62402 and GSE56814 datasets. PCSK5, ZNF225, and H1FX were used to construct a diagnostic model. ROC, calibration, and decision curve analyses were performed to assess the diagnostic performance on the training (GSE56814) and external (GSE56815) datasets. The expression level of model genes was validated in GEO datasets. Furthermore, five transcription factors (ETS1, NOTCH1, MAZ, ERG, and FLI1) were identified as common upstream regulators of model genes. PCSK5, ZNF225, and H1FX serve as novel diagnostic biomarkers, providing new insights into the pathogenesis of and treatment strategies for osteoporosis in women.

## 1. Introduction

Osteoporosis (OP) is a metabolic bone disease characterized by reduced bone mass and an impaired bone microstructure. Osteoporosis increases the risk of fracture fragility, and patients with osteoporosis frequently experience fragility fractures, with the most common sites being the lumbar spine and hip. The prevalence of OP and osteopenia was 19.7% and 40.4%, respectively, and was higher in females than in males(1). It is reported that the annual number of osteoporosis-related fractures will reach 5.99 million, and the medical cost will increase to $25.43 billion by 2050 in China(2). Hip fractures in the elderly are associated with high mortality and disability(3). With the rapid increase in the aging population, OP will become a public health problem and impose a heavy economic burden on society.

Osteoblasts and osteoclasts play critical roles in bone metabolism and are responsible for bone formation and resorption(4, 5). An imbalance in bone metabolism leads to changes in bone mineral density. Circulating monocytes play a crucial role in bone remodeling by serving as osteoclast precursors (6, 7). Monocyte subsets expressing THBS1 were found to exhibit osteoclastogenic potential through the Nucleotide-binding Oligomerization Domain (NOD)-like receptor pathway in postmenopausal osteoporosis (8). Exploring novel monocyte biomarkers using bioinformatic methods may help diagnose and treat osteoporosis earlier and broaden our understanding of OP pathogenesis. In previous studies, METTL4, RAB2A, and lncRNA ZNF529 were reported to be diagnostic biomarkers in PBMCs in postmenopausal osteoporosis(9, 10).

This study aimed to identify potential biomarkers for OP in females and develop diagnostic models for clinical practice. First, we downloaded the expression data from the public database to identify differentially expressed genes (DEGs) across two GEO datasets. GO and KEGG enrichment analyses were performed to investigate the functions of DEGs. The intersection of DEG analysis results was used to select common DEGs as biomarkers for constructing a diagnostic model. Logistic models were constructed using the “rms” package. The diagnostic efficiency was estimated using ROC, DCA, and calibration curves on the training and validation datasets. Subsequently, we validated the expression levels of the model genes in the GEO datasets. Finally, we performed transcription factor predictions and visualized them in TF-target networks. These three model genes can be regulated by five common TFs. The findings of this study identified three novel biomarkers for OP diagnosis, indicating the potential roles of PCSK5, ZNF225, and H1FX in OP pathogenesis and providing new insights and theoretical foundations for the clinical diagnosis and treatment of OP.

## 2. Materials and methods

### 2.1 Data acquisition and preprocessing

Gene expression datasets, including GSE62402, GSE56814, and GSE56815, were downloaded from the Gene Expression Omnibus (GEO; https://www.ncbi.nlm.nih.gov/geo/) database, affiliated with the National Center for Biotechnology Information (https://www.ncbi.nlm.nih.gov/). GSE62402 was performed on 5 patients with high bone density and 5 with low bone density. GSE56815 contains 40 high-BMD and 40 low-BMD samples. In the GSE56814 dataset, there were 42 high-BMD and 31 low-BMD samples. All samples were extracted from PBMCs. Detailed information about the datasets is provided in Table 1, and the workflow is shown in Fig. 1. The probe IDs were mapped to gene symbols using platform annotation files, and duplicate genes were randomly removed.

**Figure 1.**
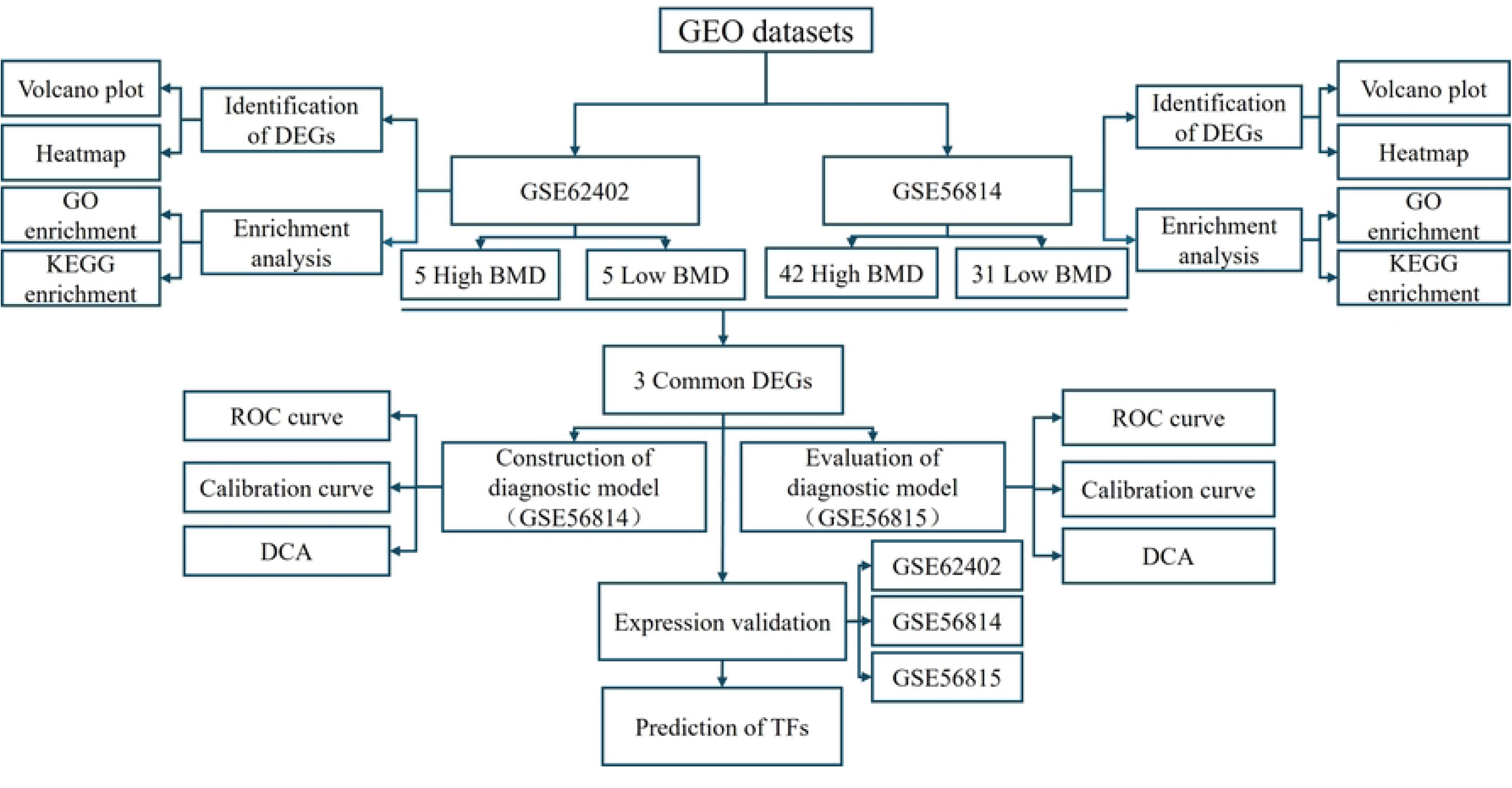
Workflow of this study. GEO = Gene Expression Omnibus, GO = Gene Ontology, KEGG = Kyoto Encyclopedia of Genes and Genomes, BMD = Bone Mineral Density, DEG = Differentially Expressed Genes, ROC = Receiver Operating Characteristic, DCA = Decision Curve Analysis, TFs = Transcription Factors.

**Table 1.**
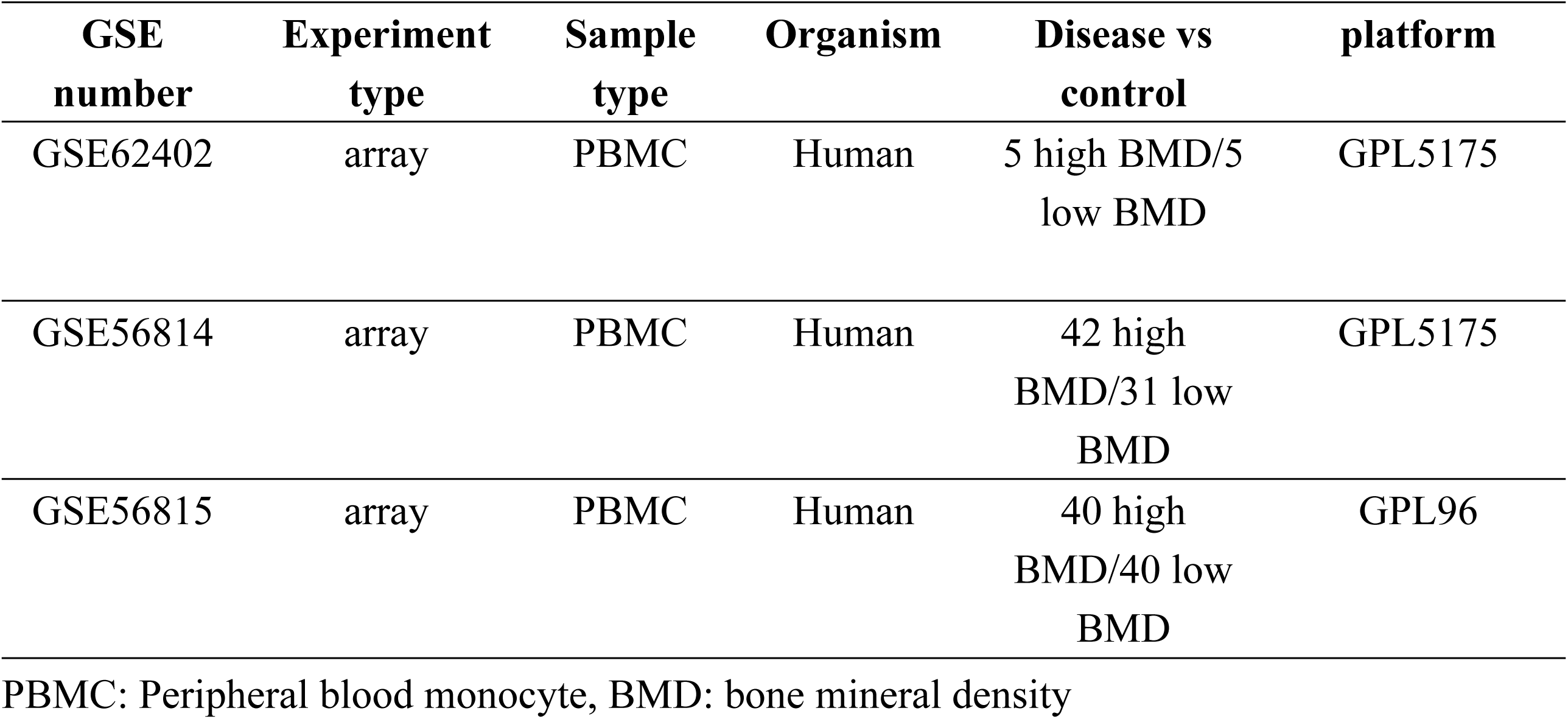
Detailed information on GEO datasets.

### 2.2 Identification and visualization of differentially expressed genes (DEGs)

We first performed principal component analysis to assess the distribution of gene expression across samples. The statistical software R and the “limma” package were used to identify DEGs with a *P*-value cutoff of <0.05 and a fold change (FC) > 1.1 (11). Heat map and volcano plot of DEGs were constructed by “ggplot2” and “pheatmap” packages.

### 2.3 GO and KEGG enrichment analyses of DEGs

GO and KEGG enrichment analyses were conducted to explore the functions of DEGs. Gene symbols were converted to Entrez IDs for enrichment analyses using the “clusterProfiler” and “org.Hs.e.g.db” packages(12). Results of GO and KEGG were visualized by the “GOplot” and “ggplot2” packages.

### 2.4 Identification of common DEGs

Based on the DEG analyses, the “VennDiagram” package was used to screen for common DEGs between GSE62402 and GSE56814.

### 2.5 Construction and evaluation of a diagnostic model for osteoporosis

We set the GSE56814 dataset as the training set, constructed diagnostic models for OP using the common DEGs, and set the GSE56815 dataset as the validation set. The “rms” package was used to build the logistic regression model. We assessed diagnostic efficiency using receiver operating characteristic (ROC) curves and the area under the curve (AUC). Decision curves were generated using the “dcurves” package. Calibration curves were plotted using the “rms” and “pROC” packages to evaluate the divergence between predicted and actual diagnostic probabilities(13).

### 2.6 Validation of model genes in GEO datasets

Gene expression profiles were obtained from the GEO datasets, GSE62402, GSE56814, and GSE56815. We applied an independent t-test to compare expression levels between the high- and low-BMD groups.

### 2.7 Prediction of transcription factors targeting the model genes

We uploaded the model genes to the hTFtarget website (https://guolab.wchscu.cn/hTFtarget/) to identify their upstream TFs and visualized the results in Cytoscape (version https://cytoscape.org/) to illustrate the common TFs that target all model genes.

## 3. Results

### 3.1 Identification of DEGs

The PCA plot shows that the samples were scattered across groups and clustered within groups (Fig. 2A-B). Using a cutoff of *P* < 0.05 and FC > 1.1, we identified 300 DEGs (247 up-regulated and 53 down-regulated) in GSE62402 and 348 DEGs (166 up-regulated and 182 down-regulated) in GSE56814. Heat maps and volcano plots were used to visualize the DEGs (Fig. 2C-F).

**Figure 2.**
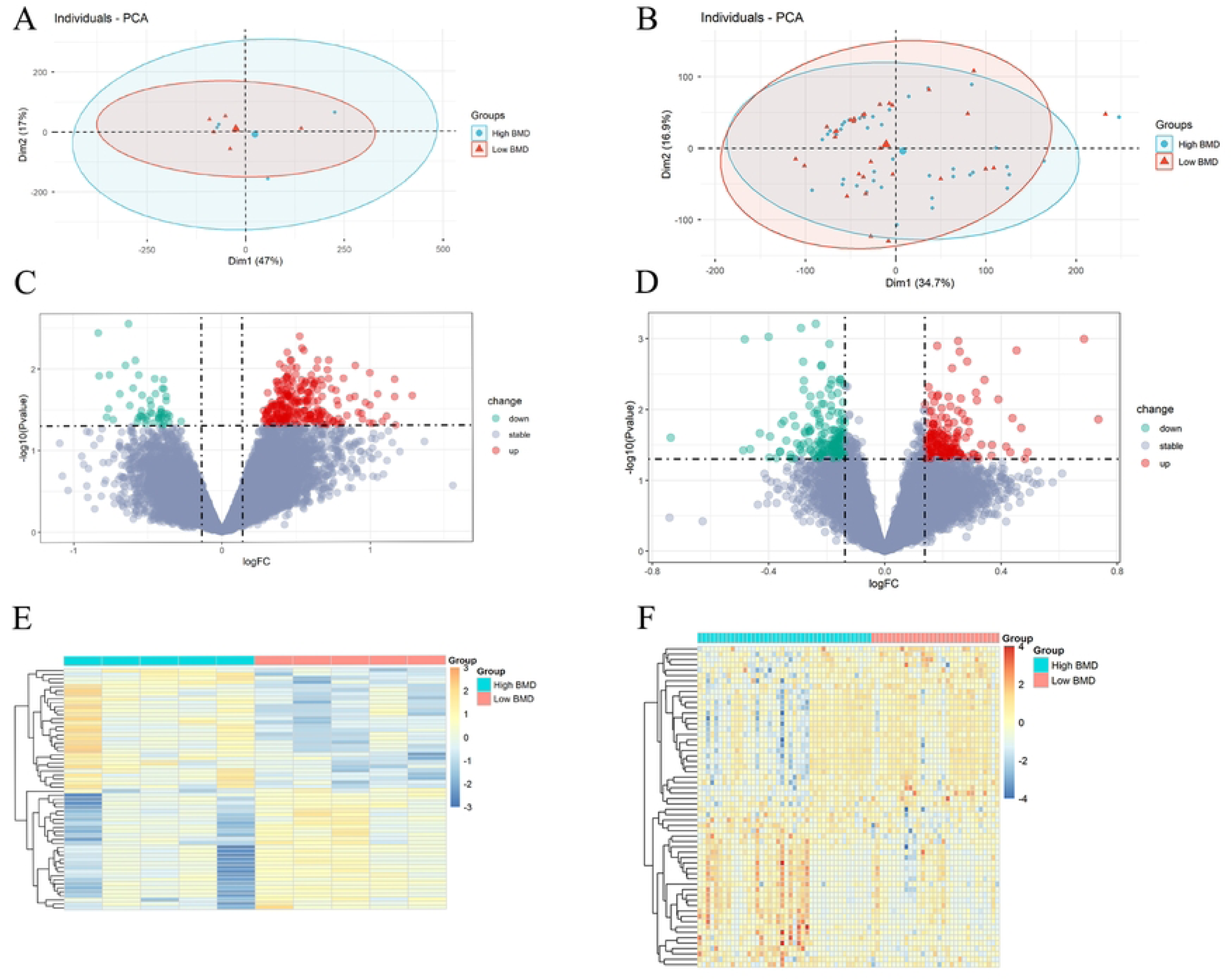
Identification of DEGs. (A-B) The principal component analysis (PCA) diagram shows the distribution of the samples (A: GSE62402; B: GSE56814). (C-D) The volcano plot of DEGs (C: GSE62402; D: GSE56814). (E-F) The heatmaps of DEGs (E: GSE62402; F: GSE56814).

### 3.2 Enrichment Analysis of DEGs

In GSE62402, GO enrichment analysis revealed the DEGs were significantly enriched in eight CCs and two MFs. CCs include COPI-coated vesicles, Golgi-associated vesicles, ficolin-1-rich granules, eukaryotic translation initiation factor 3 complexes, eukaryotic 48S preinitiation complexes, eukaryotic 43S preinitiation complexes, COPI-coated vesicle membranes, and coated vesicles. The MFs contained ribosome-binding and ribonucleoprotein complex-binding (Fig. 3A, Supplementary Table 1). As shown in Fig 3B, the top 10 KEGG pathways of DEGs were tuberculosis, lysosome, HIF-1 signaling pathway, pertussis, carbon metabolism, citrate cycle (TCA cycle), PD-L1 expression and PD-1 checkpoint pathway in cancer, protein export, Salmonella infection, and necroptosis (Supplementary Table 2). In GSE56814, GO enrichment analysis indicated that DEGs were significantly enriched in 141 BPs. The top 10 BPs were CD4-positive, alpha-beta T-cell differentiation, alpha-beta T-cell differentiation, alpha-beta T-cell activation, regulation of CD4-positive, alpha-beta T-cell differentiation, T-cell differentiation, regulation of alpha-beta T-cell activation, regulation of T-helper cell differentiation, CD4-positive, alpha-beta T-cell activation, lymphocyte differentiation, and T-cell activation involved in immune response (Fig. 3C, Supplementary Table 3). The top 10 KEGG pathways were Toll-like receptor signaling pathway, osteoclast differentiation, biosynthesis of cofactors, Kaposi sarcoma-associated herpesvirus infection, alcoholic liver disease, TNF signaling pathway, glycosphingolipid biosynthesis-lacto and neolacto series, human immunodeficiency virus 1 infection, lipid and atherosclerosis, and herpes simplex virus 1 infection (Fig. 3D, Supplementary Table 4).

**Figure 3.**
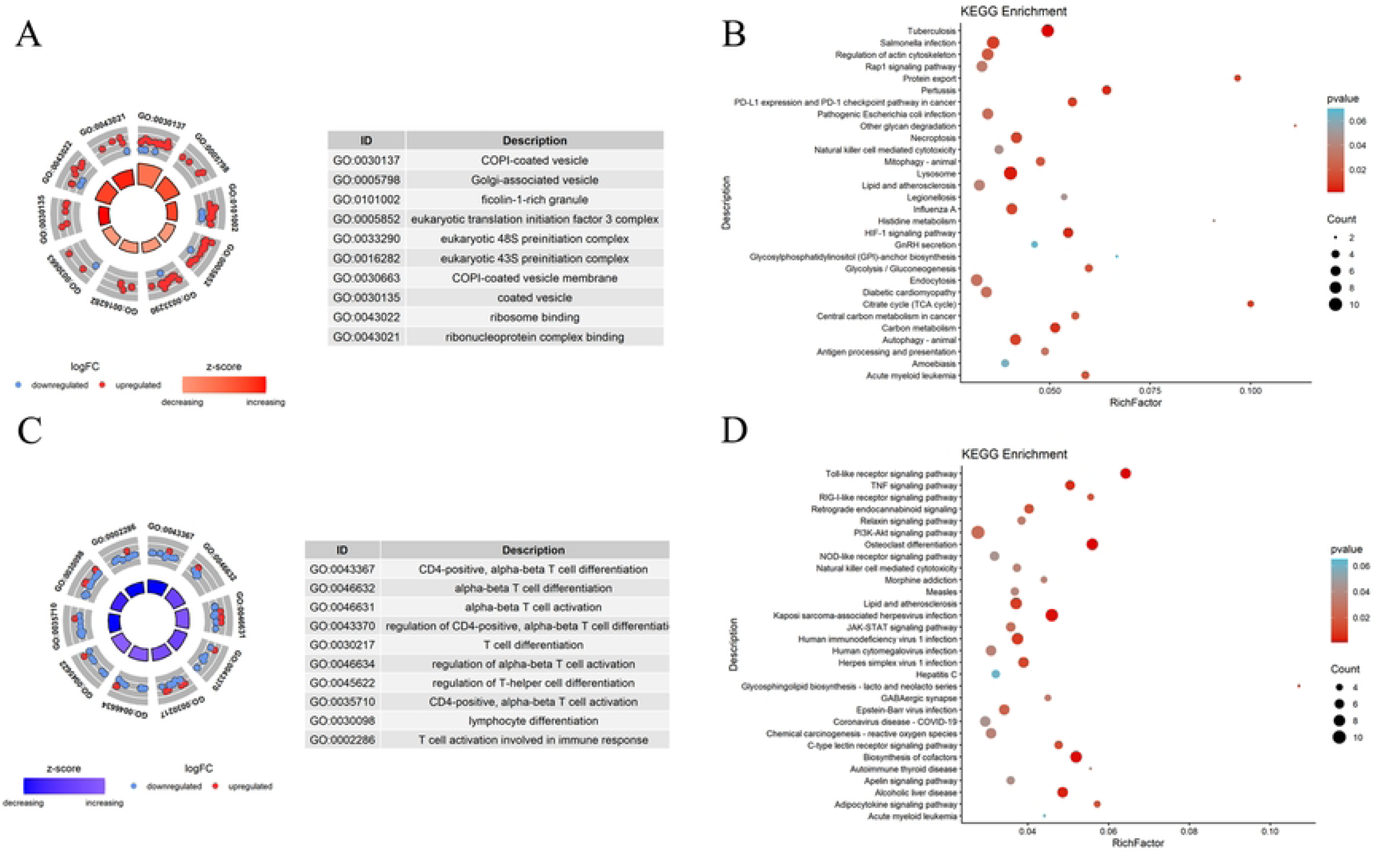
Enrichment analysis of DEGs. (A) GO enrichment analysis of DEGs in GSE62402. (B) KEGG analysis of DEGs in GSE62402. (C) GO enrichment analysis of DEGs in GSE56814. (D) KEGG analysis of DEGs in GSE56814.

### 3.3 Common DEGs screening between GSE62402 and GSE56814

As shown in Fig 4, the common upregulated genes in GSE62402 and GSE56814 were PCSK5 and ZNF225, respectively (Fig. 4A). The most commonly downregulated gene was H1FX (Fig. 4B).

**Figure 4.**
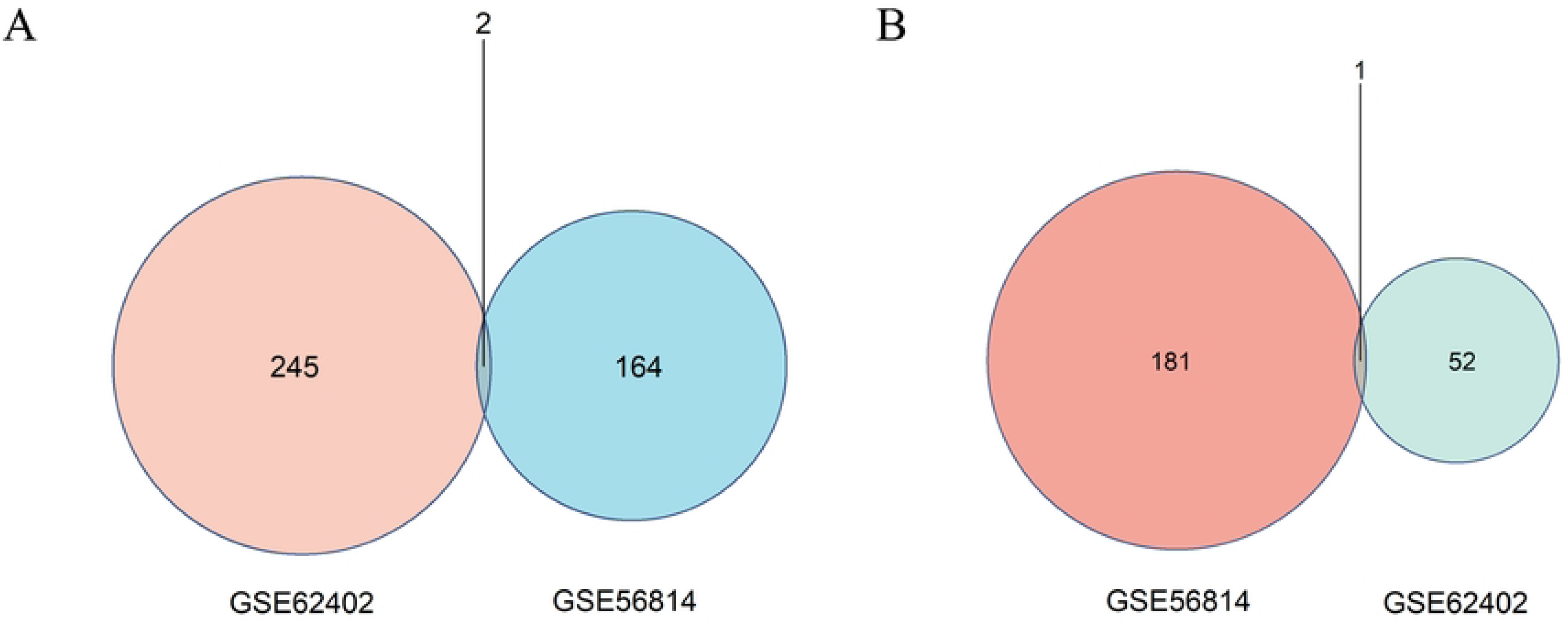
Identification of common DEGs. (A) The Venn diagram of commonly upregulated DEGs in the GSE62402 and GSE56814 datasets. (B)The Venn diagram of commonly downregulated DEGs in the GSE62402 and GSE56814 datasets.

### 3.4 Diagnostic Model for OP

Common DEGs, including PCSK5, ZNF225, and H1FX, were selected as biomarkers for constructing the OP model. The diagnostic models were constructed by the “rms” R package in the GSE56814 dataset. As shown in Fig. 5A, the AUCs for PCSK5, ZNF225, and H1FX were 0.678, 0.652, and 0.636, respectively. The *P* values for PCSK5, ZNF225, and H1FX in the diagnostic model calibration curve were all greater than 0.05, indicating good consistency and diagnostic efficiency of the model (Fig. 5B-D). The DCA plot revealed that all three models could benefit patients in clinical diagnosis (Fig. 5E). In the validation dataset (GSE56815), the AUCs for PCSK5, ZNF225, and H1FX were 0.851, 0.876, and 0.793, respectively (Fig. 6A). The *P* values for PCSK5 and H1FX in the diagnostic model calibration curve were all greater than 0.05, whereas the P value for ZNF225 was less than 0.05 (Fig. 6B-D). DCA analysis indicated all models were excellent for the diagnosis of OP (Fig. 6E).

**Figure 5.**
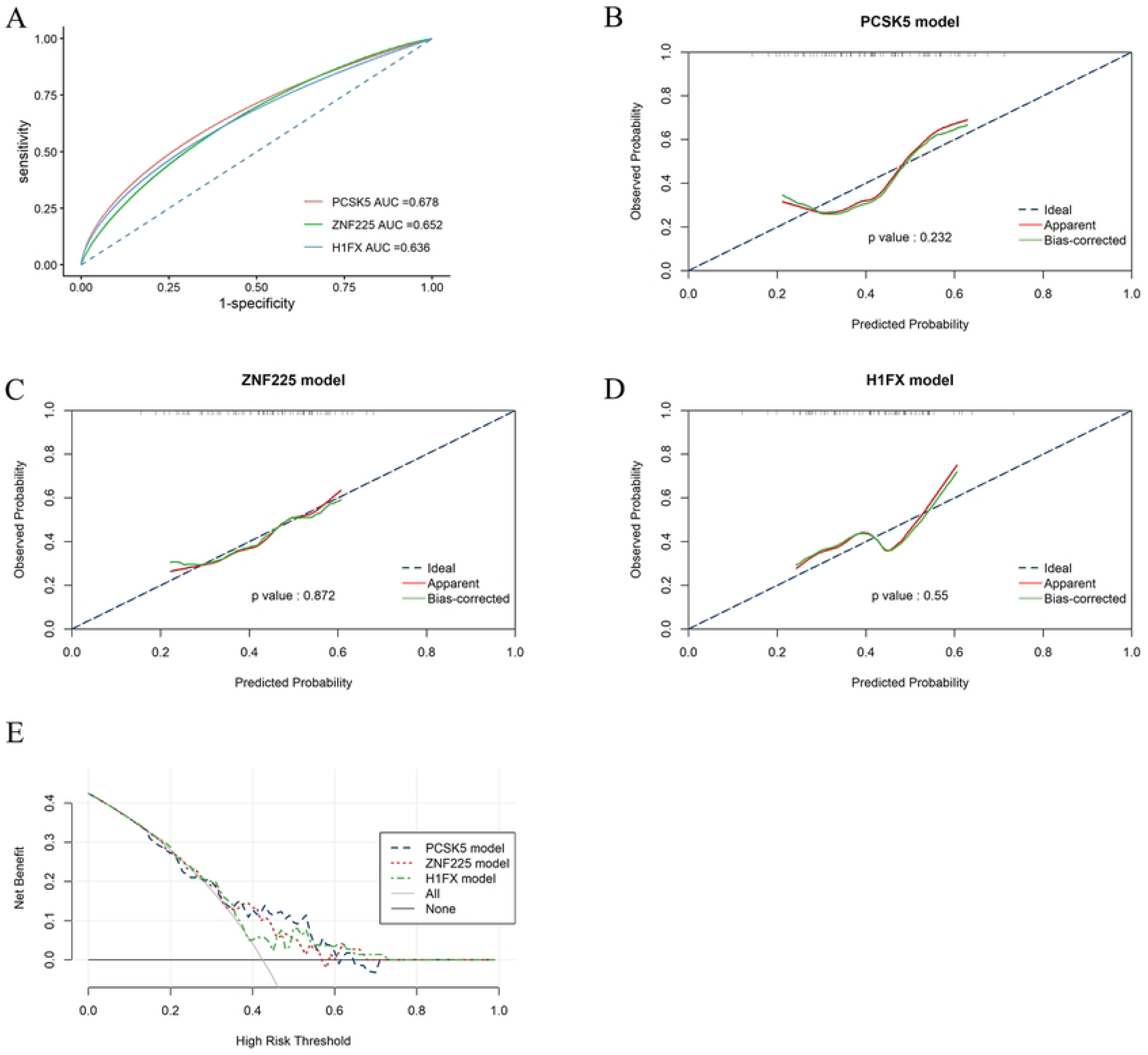
The establishment of diagnostic models for OP in the GSE56814 dataset. (A) ROC analysis of diagnostic models based on PCSK5, ZNF225, and H1FX in the training dataset. (B-D) Calibration curve of diagnostic models for PCSK5, ZNF225, and H1FX (B: PCSK5; C: ZNF225; D: H1FX) (*P* > 0.05). (E) DCA curves of PCSK5, ZNF225, and H1FX models.

**Figure 6.**
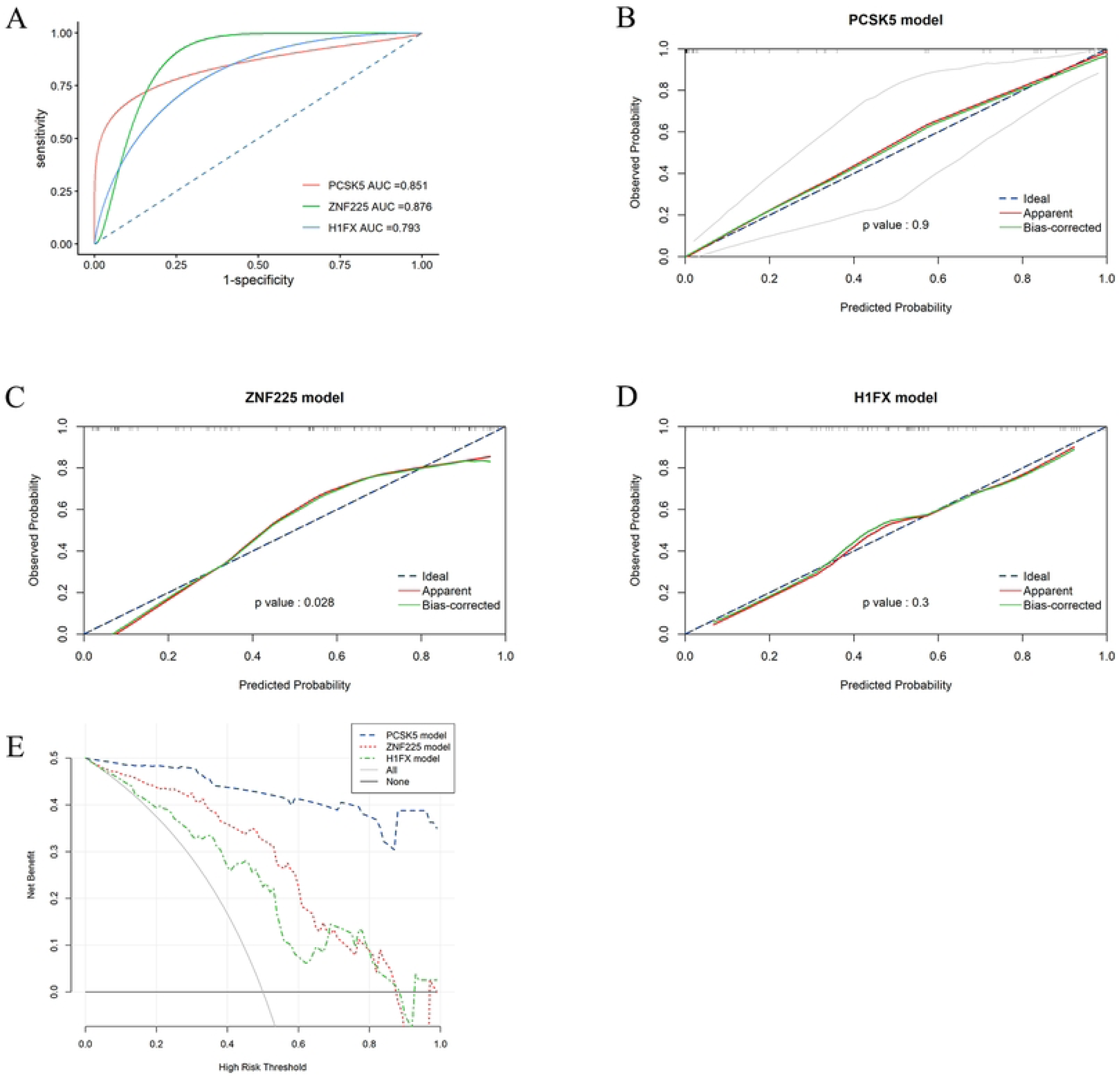
The evaluation of diagnostic models for OP in the GSE56815 dataset. (A) ROC analysis of diagnostic models based on PCSK5, ZNF225, and H1FX in the training dataset. (B-D) Calibration curve of diagnostic models for PCSK5 (*P* > 0.05, ZNF225 (*P* < 0.05, and H1FX (*P* > 0.05) (B: PCSK5; C: ZNF225; D: H1FX). (E) DCA curves of PCSK5, ZNF225, and H1FX models.

### 3.5 Expression validation of model genes in the datasets

We further validated the expression of key genes in the datasets, including GSE62402, GSE56814, and GSE56815. Consistently, the expression of ZNF225 and PCSK5 was significantly higher in the low BMD group than in the high BMD group, as assessed by Student’s t-test (*P* < 0.05) (Fig. 7A-C). The expression of H1FX was significantly lower in the low BMD group than in the high BMD group (P < 0.05) (Fig. 7A-C).

**Figure 7.**
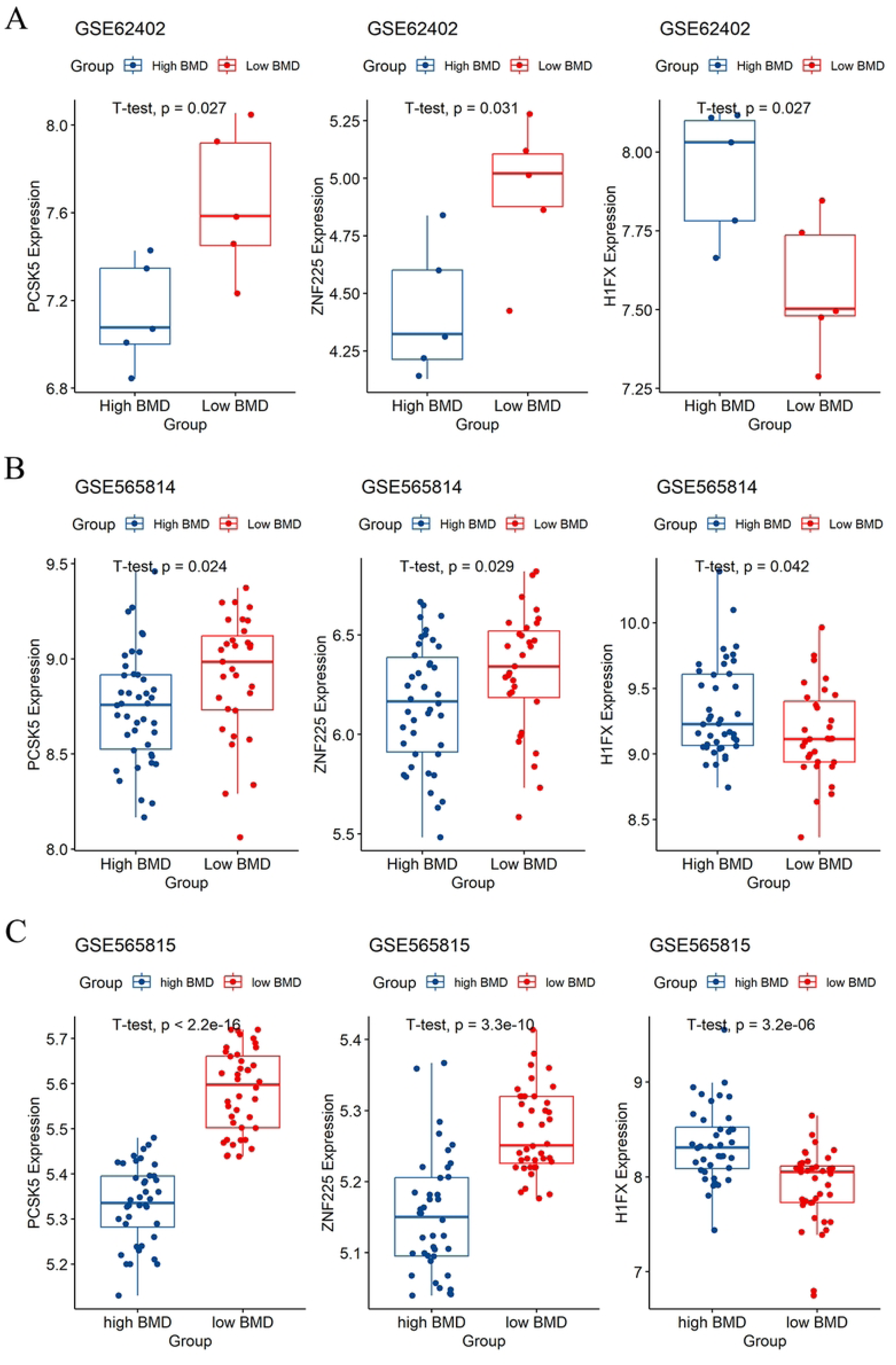
Expression validation of biomarkers in GEO datasets. (A) The expression comparison of PCSK5, ZNF225, and H1FX between the high-BMD and low-BMD groups in GSE62402 (t-test, *P* < 0.05). (B) The expression comparison of PCSK5, ZNF225, and H1FX between the high-BMD and low-BMD groups in GSE56814 (t-test, *P* < 0.05). (B) The expression comparison of PCSK5, ZNF225, and H1FX between the high-BMD and low-BMD groups in GSE56815 (t-test, *P* < 0.05).

### 3.6 Prediction of transcription factors targeting the key genes

We constructed predictions for key genes on the hTFtarget websites. Consequently, 22 TFs were predicted to modulate ZNF225 expression, and 13 TFs were predicted to be regulators of PCSK5. 69 TFs were selected as upstream factors of H1FX (Supplementary Table 5). In addition, three key genes can be regulated by five TFs, including ETS1, NOTCH1, MAZ, ERG, and FLI1(Fig. 8A-B).

**Figure 8.**
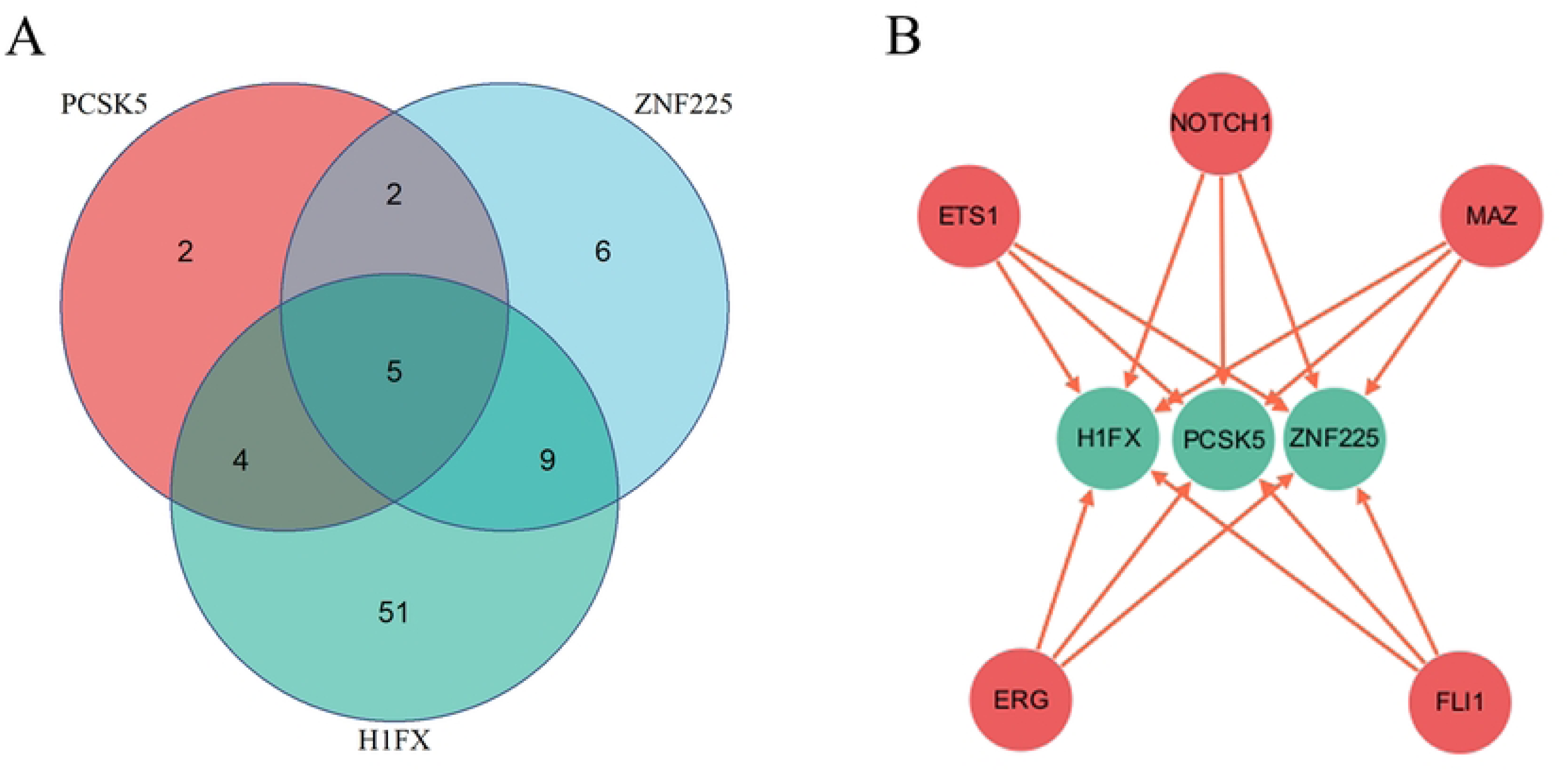
Identification of common upstream TFs among PCSK5, ZNF225, and H1FX. (A) Venn diagram of common upstream TFs among PCSK5, ZNF225, and H1FX. (B) The TF-target network of common TFs and biomarkers.

## 4. Discussion

Osteoporosis is a systemic metabolic skeletal disorder characterized by low BMD and deterioration of bone microstructure(14). The prevalence of osteopenia and OP was high. In the latest report in the United States, 13%-18% of women aged 50 years had osteoporosis, and 37%-50% had osteopenia(15). In addition, approximately 200 million women have osteoporosis. The prevalence of OP in women has been reported to be rising. Patients with OP often suffer from a fragility fracture in the vertebrae and hip, and the prognosis is poor. In the US, the cost of treating one case of fragility fracture ranges from $5,000 to $19,000 (16). Dual-energy X-ray absorptiometry (DXA) is an X-ray-based technique that quantifies bone mineral density and is widely used in the diagnosis of OP. Osteoporosis was defined as a T-score below -2.5(17). Current treatments for OP usually target bone formation or bone resorption. PBMCs play a role in bone mass and serve as precursors to regulate osteoclast differentiation and activity(18).

In our study, we used bioinformatics analysis to screen for novel biomarkers for OP diagnosis. First, 300 and 348 DEGs were identified between the low- and high-BMD groups in GSE62402 and GSE56814, respectively. As shown in the GO enrichment analysis results, the DEGs in GSE62402 were significantly enriched in 8 CCs and 2 MFs, whereas the DEGs in 56814 were considerably enriched in 141 BPs. Further KEGG analysis revealed that the DEGs in GSE62402 were mainly enriched in tuberculosis, lysosome, HIF-1 signaling pathway, pertussis, carbon metabolism, citrate cycle (TCA cycle), PD-L1 expression and PD-1 checkpoint pathway in cancer, protein export, Salmonella infection, and necroptosis. The top 10 KEGG pathways in GSE56814 analysis were Toll-like receptor signaling pathway, osteoclast differentiation, biosynthesis of cofactors, Kaposi sarcoma-associated herpesvirus infection, alcoholic liver disease, TNF signaling pathway, glycosphingolipid biosynthesis-lacto and neolacto series, human immunodeficiency virus 1 infection, lipid and atherosclerosis, and herpes simplex virus 1 infection. Two upregulated DEGs (PCSK5 and ZNF225) and one downregulated gene (H1FX) were selected as common DEGs to construct the OP diagnostic model. In the training dataset, the AUCs for PCSK5, ZNF225, and H1FX were 0.678, 0.652, and 0.636, respectively. As in the external validation dataset, the AUCs for PCSK5, ZNF225, and H1FX were 0.851, 0.876, and 0.793, respectively. DCA and calibration curves for both the training and validation datasets demonstrated excellent diagnostic efficiency. Consistently, the expression of PCSK5 and ZNF225 was significantly up-regulated, and that of H1FX was significantly down-regulated, in the low-BMD group in GSE62402, GSE56814, and GSE56815. Moreover, PCSK5, ZNF225, and H1FX were shown to be regulated by five common TFs: ETS1, NOTCH1, MAZ, ERG, and FLI1. Zinc finger proteins have been reported to play important roles in many biological processes, including hepatocellular carcinoma (HCC) progression and hematopoiesis (19, 20). Choi reported that the combination of everolimus and Ku0063794 exerts an anticancer effect by reducing cell autophagy through the upregulation of ZNF225 (21). In another comprehensive analysis, ZNF family genes, including ZNF225, were strongly associated with prognosis in esophageal cancer(22). However, the exact function of ZNF225 in the skeletal system has not yet been described.

PCSK5 (Proprotein Convertase Subtilisin/Kexin Type 5) is a protein-precursor convertase responsible for proprotein processing. In previous studies, PCSK5 was expressed in osteoblasts and osteocytes, and proved to be indispensable for skeletal development(23, 24). It is upregulated during the osteogenic differentiation of human bone marrow-derived mesenchymal stromal cells (BMSCs) in vitro and during the bone-healing period of murine tibial fracture in vivo(25). Furthermore, miR-338-3p was shown to promote age-associated osteoporosis by inhibiting osteogenesis through downregulation of PCSK5 expression in an in vitro study (26). In addition, PCSK5 has been identified as a biomarker for endometriosis and inflammatory skin diseases (27, 28).

As reported by Liang et al., H1FX has been screened as a biomarker for allergic rhinitis with excellent diagnostic efficiency(29). In proteomic analysis, H1FX was identified as a biomarker for stratifying prostate cancer from low- and high-grade groups(30). In a recent single-cell RNA sequencing study, H1FX was shown to regulate the Neut_RSAD2 subpopulation, which is involved in the bone microenvironment in osteoporosis (31). ETS1, NOTCH1, MAZ, ERG, and FLI1 were predicted as upstream TFs for the three model genes and may be potential targets for OP.

This study revealed that PCSK5, ZNF225, and H1FX were closely associated with osteoporosis in women. Our study has several limitations. First, all analyses were performed using public datasets with bioinformatics methods. Second, exact bone mineral density data were not available in the datasets, so we could not carry out a correlation analysis between the key genes and BMD. Third, we did not validate the expression of the hub genes in vivo. In the future, we will further confirm the potential role of these three genes in osteoclastogenesis.

## Data Availability

Our data can be found in the Gene Expression Omnibus (GEO; https://www.ncbi.nlm.nih.gov/geo/) database under accession numbers GSE62402, GSE56814, and GSE56815.

## Authors’ contributions

Conception and design: X.Q and Z.M; Data collection and analysis: B.W., P.W., Z.C., and Z.M; Manuscript writing and revision: X.Q., B.W., and S.T. All authors read and approved the final manuscript.

## Acknowledgements

Not applicable.

## Abbreviations

OP: osteoporosis
GEO: Gene Expression Omnibus
DEG: Differentially Expressed Genes
GO: Genome Ontology
KEGG: Kyoto Encyclopedia of Genes and Genomes
AUC: Area under the receiver operating characteristic curve
FC: Fold Change
PCA: Principal component analysis
BP: Biological processes
CC: Cellular component
MF: Molecular function
PCSK5: Proprotein Convertase Subtilisin/Kexin Type 5
ZNF225: Zinc Finger Protein 225
H1FX: H1 Histone Family Member X
TF: transcription factor
DCA: decision curve analysis
ROC: receiver operating characteristic
BMD: bone mineral density
DXA: Dual-energy X-ray absorptiometry
PBMCs: peripheral blood mononuclear cells
HCC: hepatocellular carcinoma

